# Comparative analysis of mouse strains for *in vivo* reprogramming

**DOI:** 10.1101/2024.03.08.584074

**Authors:** Sara Picó, Alba Vílchez-Acosta, João Agostinho de Sousa, María del Carmen Maza, Alberto Parras, Gabriela Desdín-Micó, Calida Mrabti, Céline Yacoub Maroun, Clémence Branchina, Ferdinand von Meyenn, Alejandro Ocampo

## Abstract

*In vivo* reprogramming through the forced expression of Oct4, Sox2, Klf4, and c-Myc (OSKM) has demonstrated great potential for reversing age-associated phenotypes, as the combination of these transcription factors actively promote cell regeneration and rejuvenation in various tissues and organs. However, continuous *in vivo* OSKM expression raised safety concerns due to loss of cell identity, decrease in body weight, and premature death. Although cyclic short-term or targeted expression of the reprogramming factors can mitigate some of these detrimental effects in mice, systemic rejuvenation of wild type mice has remained elusive potentially due to these current technical limitations. To improve the fundamental understanding of *in vivo* reprogramming, we conducted a comparative analysis across multiple reprogrammable mouse strains, tissues, and expression methods, presenting a comprehensive atlas of formerly established strains. In addition, we developed novel reprogrammable mouse strains by avoiding OSKM expression in specific organs, in dividing cells, or implementing chimeric expression approaches within specific cells, thereby offering safer strategies to induce *in vivo* reprogramming and fully harness its potential. We hope that these new tools will become valuable resources for future research in this very exciting field of research with potential implications to human health.

## Introduction

In 2006, Shinya Yamanaka demonstrated that the expression of the four transcription factors *Oct4*, *Sox2*, *Klf4* and *c-Myc* (OSKM), is capable of reprogramming differentiated cells in vitro back to a pluripotent state^1^. Intriguingly, subsequent studies have shown that induction of partial reprogramming can reverse age-associated phenotypes in vitro, highlighting its potential to modify both cellular identity and age^2–4^. In this line, OSK(M) have been shown to promote the regeneration of multiple cell types and restore young transcription profiles^5^. Furthermore, tissues and organs that have been rejuvenated by *in vivo* reprogramming include kidney^6^, liver^7,8^, skin^6,9^, heart^10^, pancreas and muscle^4,7,11–13^. In addition, brain memory ^14^ and axon regeneration after injury^15^ have been improved by *in vivo* reprogramming.

Despite the potential opportunities proposed by advances in cellular reprogramming, the continuous induction of OSKM expression *in vivo* faces notable safety issues because of the loss of cell identity^16–18^ leading to organ failure and dysfunction, severe body weight loss, and early mortality^8,11,16,19^. Importantly, these defects can be uncoupled from the rejuvenating effects of *in vivo* reprogramming by controlling the expression of reprogramming factors, either by cyclic short-term expression or cell-or tissue-specific expression. Importantly, partial reprogramming, achieved by transient periodic induction of OSKM *in vivo*, ameliorated signs of aging without loss of cellular identity in progeroid mice, extending their lifespan in the absence of teratoma formation^11^. However, it is evident that these methods, whether short-term or mildly induced, have failed to induce systemic rejuvenation or extend the lifespan of wild-type mice. In this line, a recent study using gene therapy to induce partial reprogramming showed a mild effect on organismal rejuvenation and lifespan extension in physiologically aged mice^20^. Hence, we are in need of developing novel strategies that can facilitate the safe induction of *in vivo* reprogramming, minimizing its adverse side effects, to fully realize its true rejuvenation potential.

Towards this goal, we conducted a comparative analysis of *in vivo* reprogramming at both phenotypic and OSKM expression levels in previously established reprogrammable mouse strains across different tissues. First, we analyzed four different strains in the context of whole-body reprogramming. Additionally, we analyzed the potential effects of changes in the number of copies of the OSKM cassette and transactivator, as well as the differences in induction protocols. Lastly, with the goal of establishing new protocols for the induction of safer, stronger, and long-term *in vivo* reprogramming, we generated and analyze novel reprogrammable mouse strains based on OSKM expression in specific tissues (avoiding expression in the liver and intestine), expression of the reprogramming factors in post-mitotic cells, or expression in a chimeric fashion that allows to sustain organ function during *in vivo* reprogramming. We strongly believe that this study provides a deep perspective of *in vivo* reprogramming, offering valuable data for the study of safe reprogramming strategies to induce rejuvenation at the organismal level.

## Results

### Transcriptome analysis unveils differential expression of the reprogramming factors between different reprogrammable mouse strains

With the aim of establishing better strategies for the induction of whole-body *in vivo* reprogramming, we decided to explore the expression of OSKM in four previously established reprogrammable mouse strains. Importantly, the main genetic difference between these strains is the genetic location of the doxycycline-inducible polycistronic cassette (TetO 4F), which encodes murine *Oct4*, *Sox2*, *Klf4,* and *c-Myc* (OSKM). While in the 4Fj and 4Fk strains, the TetO 4F is inserted in the *Col1a1* locus, in the 4FsB is in the *Ppar*g locus and in the 4FsA in the *Neto2* locus. At the same time, the order of Yamanaka factors in the polycistronic cassette is OSKM for 4Fj, 4FsB, 4FsA, while it is OKSM in the case of the 4Fk strain (Figure 1A, left).

**Figure 1.**
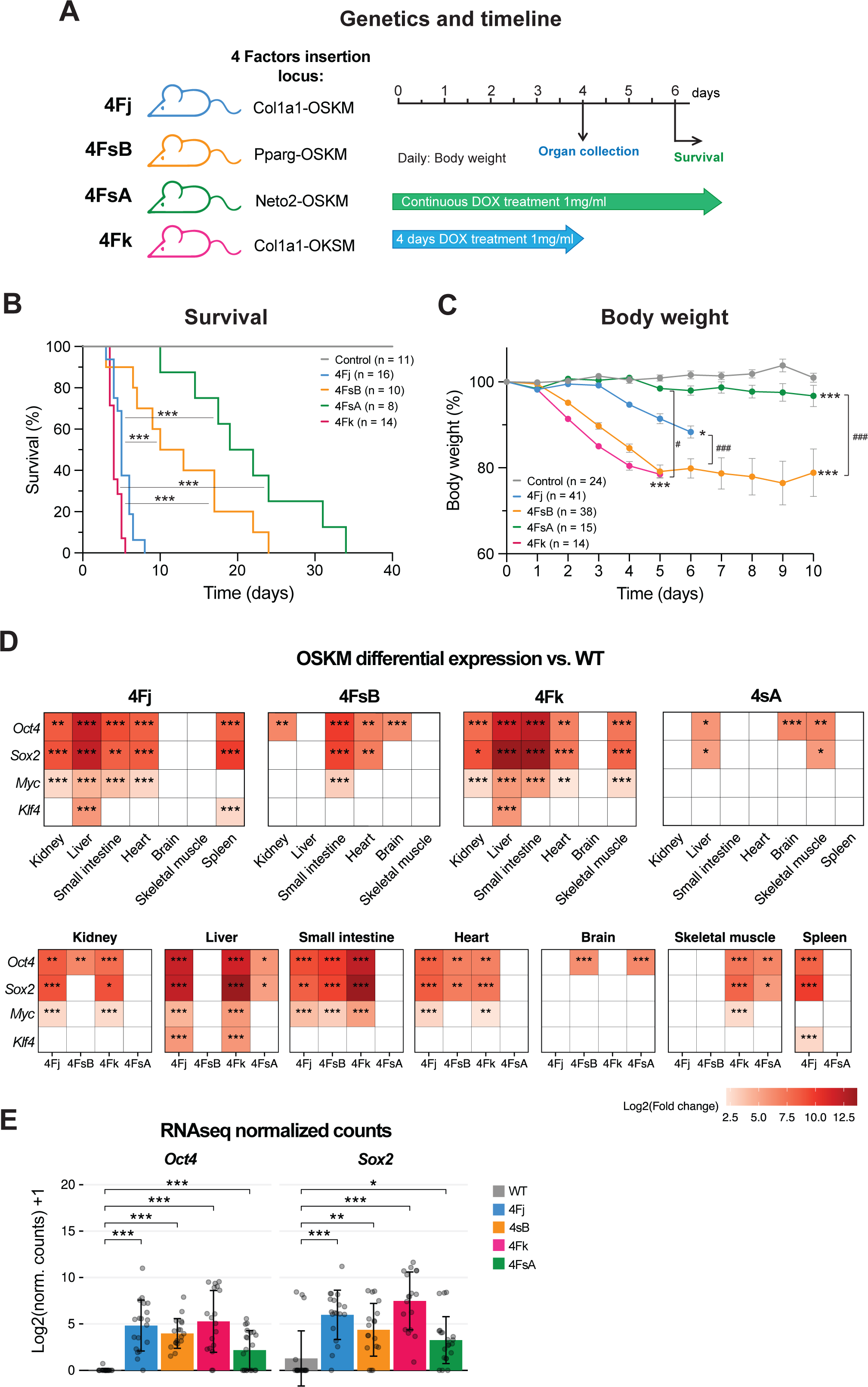
Comparative *in vivo* reprogramming in different whole body reprogrammable mouse strains. **A**, Graphical representation of reprogrammable mouse strains carrying the polycistronic cassette for the reprogrammable factors in different loci (Col1a1, Pparg and Neto2) and with different order of the Yamanaka factors (OSKM or OKSM). **B,** Survival of 4Fj, 4FsB, 4FsA and 4Fk upon continuous administration of doxycycline and control (WT) mice. **C,** Body weight changes in 4Fj, 4FsB, 4FsA and 4Fk upon continuous administration of doxycycline and in control (WT) mice. **D**, Differential expression analysis comparing the 4F strains to the wild-type strain for each OSKM gene across different tissues after 4 days of doxycycline treatment. Positive Log2 Fold change means the WT samples have lower expression. **E,** RNA-seq log-transformed normalized counts of *Oct4* and *Sox2* genes across mouse strains in all tissues. Data shown mean ± standard deviation. Statistical significance was assessed by one-way ANOVA followed by Tukey’s post hoc test (c), R bioconductor package DESeq2 (d), T-test (E) and log-rank (Mantel–Cox) test (b). See also Figure S1.

First, we induced the expression of OSKM in all strains at 2 months of age by continuous administration of doxycycline (1 mg/ml) in drinking water and monitor changes in body weight and survival (Figure 1a). As expected, the 4Fj and 4Fk strains, in which the transgene is in the *Col1a1* locus, showed reduced median survival compared to the 4FsA and 4FsB strains (Figure 1B). In addition, a decrease in body weight was observed in all strains to a different extent upon induction of the reprogramming factors compared to controls (Figure 1C).

To investigate whether these results were correlated with different levels of expression of OSKM, we performed global transcriptomic analysis using RNA sequencing (RNA-seq) across several tissues and organs after 4 days of doxycycline administration (1 mg/ml) in drinking water (Figure 1A, right). Principal component analysis (PCA) revealed a clear separation among tissue types in different clusters and importantly, revealed a clear separation between reprogrammable and control strains within each tissue (Figure S1A). As expected, liver and small intestine were the organs with the most significant increase in transcript levels of OSKM in 4Fj and 4Fk strains that show early mortality, followed by kidney (Figure 1D). However, in organs unrelated to doxycycline absorption, such as heart and skeletal muscle, OSKM transcript levels were more similar between the strains (Figure 1D). Similar results were obtained by qPCR (Figure S1B). In addition, 4Fk and 4Fj strains express higher levels of the reprogramming factors in average in all the tissues and organs analyzed (Figure 1E), correlating with their reduced survival.

Overall, these results suggest that OSKM expression is different across various reprogrammable mouse strains and higher in tissues and organs that are more exposed to doxycycline such as liver, intestine, and kidney, particularly in the 4Fj and 4Fk strains.

### OSKM expression alters global transcriptome

To capture the possible transcriptional heterogeneity generated during *in vivo* reprogramming, we next analyzed RNA-seq data in more depth. After comparing differentially expressed genes (DEGs) in all tissues, we observed that the 4Fk strain exhibited a higher number of DEGs, in agreement with the enhanced expression of OSKM factors in this strain (Figure 2A). Interestingly, when we explored DEGs for each tissue, we found a correlation of DEGs levels and OSKM expression mainly in liver and intestine of 4Fk, however, in tissues like the 4sA heart and spleen, DEGs were still present despite the minimal OSKM expression, suggesting that the expression in other tissues might have organ-extrinsic effects on DEGs (Figure 2A). In addition, our Gene ontology (GO) analysis showed that upregulated DEGs were significantly enriched in terms related to response to inflammation, infections, and regulation of blood clotting, while downregulated DEGs were enriched in terms related to immune responses, including both the innate and adaptive immune systems (Figure 2B). In conclusion, these results demonstrate that whole-body *in vivo* reprogramming alters the global transcriptome in a direct or indirect manner, resulting in differentially expressed genes even in tissues with low OSKM expression.

**Figure 2.**
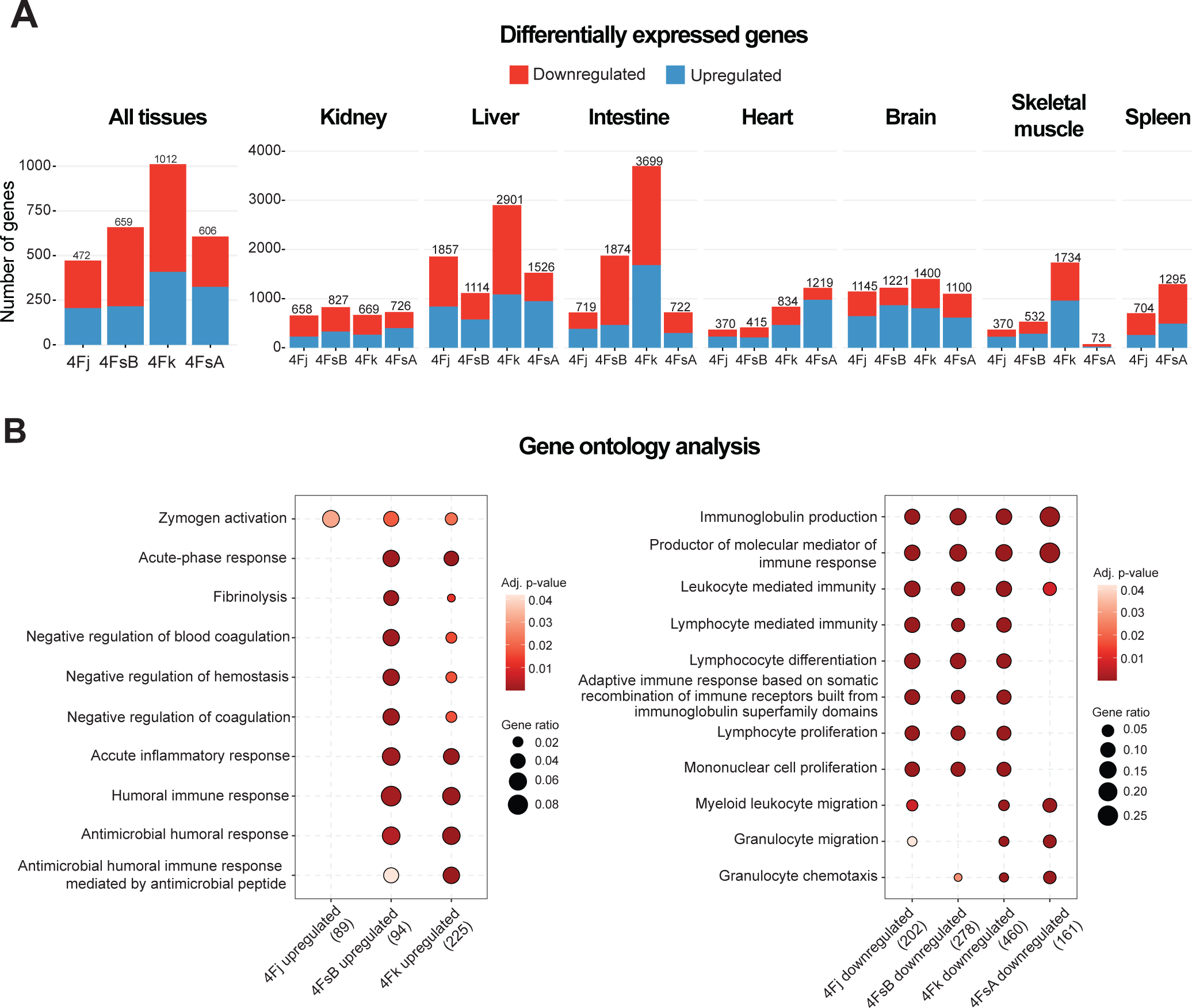
Global transcriptome changes in reprogrammed strains. **A**, Differentially expressed genes (DEGs) in all tissues (left) and separated by tissue (right) of 4Fj, 4FsB, 4FsA and 4Fk vs. control (WT) mice. **B,** Gene ontology (GO) analysis of DEGs in all tissues of 4Fj, 4FsB, 4FsA and 4Fk vs. control (WT) mice.

### Expression levels of OSKM can be controlled by copy number of the reprogramming cassette

Given that the 4Fj mouse strain showed one of the highest expressions of OSKM factors across the analyzed tissues, we explored how different genetic strategies regarding the number of copies of the TetO 4F and reverse tetracycline-controlled transactivator (rtTA) cassettes could modulate the mRNA levels of OSKM in this strain. First, we compared side by side the effects of mouse carrying two copies of the TetO 4F cassette (4FjF rtTA HOMO) or the replacement of rtTA by the new-generation rtTA3 (4FjF rtTA3 HET)^21^ (Figure 3A). Upon induction of reprogramming by administration of 1mg/ml doxycycline in drinking water to 2 months old mice, a significant reduction in survival and body weight loss was observed in the two newly generated 4FjF rtTA HOMO and 4FjF rtTA3 HET mice compared to 4FjF rtTA HET mice (Figure 3B and Figure S2A). In addition, we studied the expression levels of the factors in several tissues, after 3-4 days of doxycycline treatment, and found a significant increase in *Oct4* and *Sox2* transcript levels in multiple tissues and organs of 4FjF rtTA HOMO mice and, to a milder extent, in 4FjF rtTA3 HET mice compared to 4FjF rtTA HET mice (Figure 3C and Figure S2B). Next, we explored whether increasing the concentrations of doxycycline could modulate the expression levels of these factors in different tissues. To this end, three different concentrations of doxycycline (1, 2, and 5 mg/ml) were given to the 4FjF rtTA HET mouse at 2 months. Animals treated with the higher doses (2 and 5 mg/ml) showed similar decrease in survival and body weight loss compared to mice treated with 1 mg/ml of doxycycline (Figure 3D-E). However, these phenotypic differences did not correlate with the levels of *Oct4* and *Sox2* transcripts in the tissues analyzed, which were not significantly different at different doxycycline concentrations (Figure 3F and Figure S2C).

**Figure 3.**
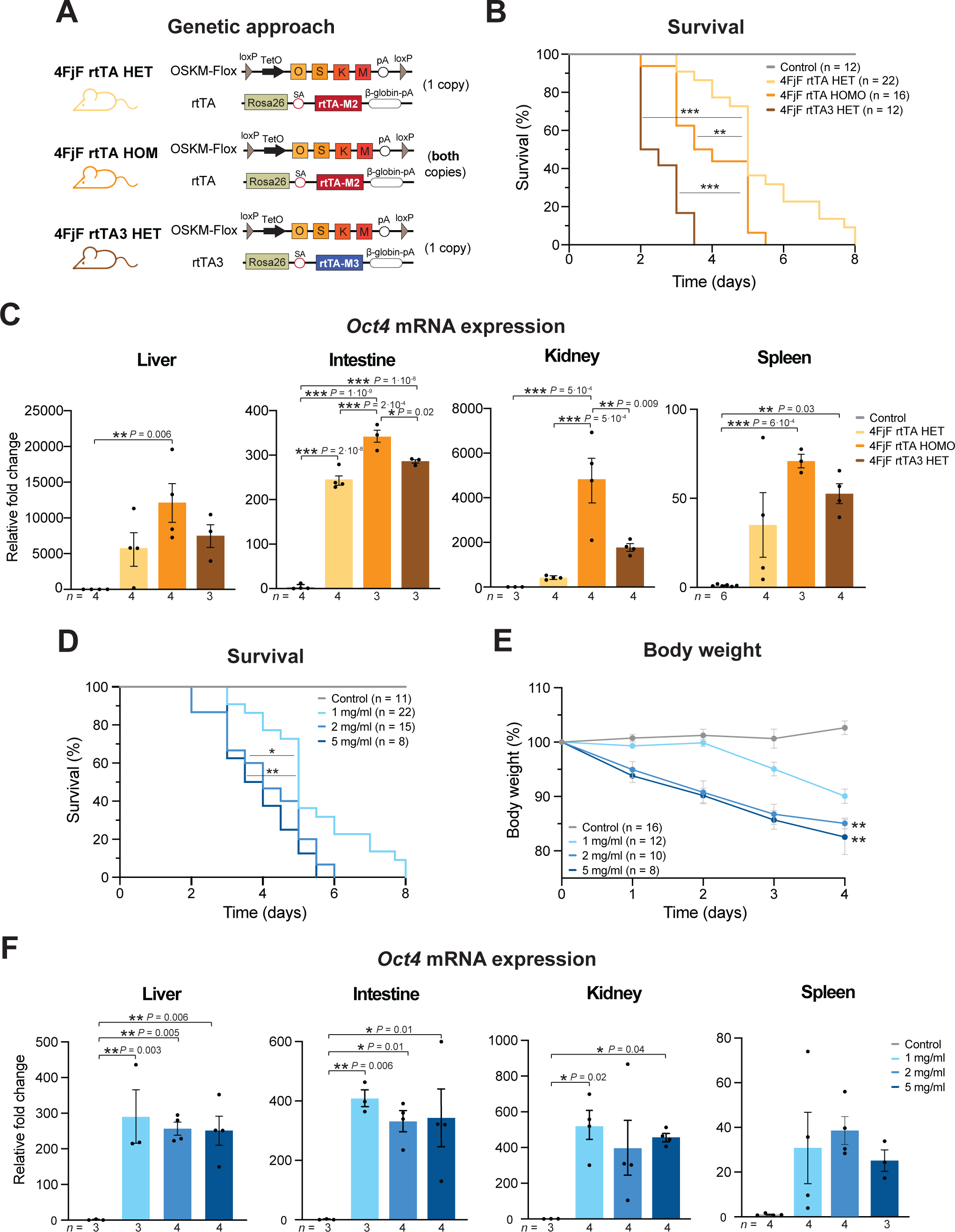
Strategies to increase the levels of *in vivo* reprogramming. **A**, Schematic representation of the genetic approaches used. The 4FjF rtTA mouse strain, carrying the reverse tetracycline-controlled transactivator transgene, rtTA-M2, at the Rosa26 locus and one (HET) or two (HOMO) copies of the inducible polycistronic cassette for the expression of the murine Yamanaka factors (*Oct4*, *Sox2*, *Klf4* and *cMyc*), between two loxP sites, OSKM-Flox, (pA) polyA sequence, (TetO) tetracycline operator minimal promoter. The 4FjF rtTA3 in which the doxycycline transactivator is replaced by the rtTA-M3 transgene. **B,** Survival of control, 4FjF rtTA HET, 4FjF rtTA HOMO and 4FjF rtTA3 HET mice upon continuous administration of doxycycline. **C,** *Oct4* mRNA transcript levels in multiple organs in control (WT), 4FjF rtTA HET, 4FjF rtTA HOMO and 4FjF rtTA3 HET after 3-4 days of doxycycline treatment. **D,** Survival of control (WT) and 4FjF rtTA HET mice upon continuous administration of doxycycline at 1mg/ml 2 mg/ml and 5mg/ml. **e,** Body weight changes in control (WT) and 4FjF rtTA HET mice upon continuous administration of doxycycline at 1mg/ml 2 mg/ml and 5mg/ml. **f,** *Oct4* mRNA transcript levels in multiple organs in control (WT) and 4FjF rtTA HET after 4 days of doxycycline treatment at 1mg/ml 2 mg/ml and 5mg/ml. Data shown mean ± standard deviation. Statistical significance was assessed by one-way ANOVA followed by Tukey’s post hoc test (c,e-f) and log-rank (Mantel–Cox) test (b,d). See also Figure S2.

Taken together, these results demonstrate that both genetic strategies increasing cassette copy number, using a more powerful transactivator, or increasing the concentration of doxycycline significantly enhance the expression of OSKM during reprogramming and impact body weight and survival.

### Enhanced *in vivo* reprogramming efficiency by avoiding OSKM expression in liver and intestine

A previous study by our group has recently shown that *in vivo* reprogramming leads to hepatic and intestinal dysfunction and represents one of the major causes of loss of body weight and early mortality upon induction of OSKM expression. Consequently, bypassing the expression of OSKM in the liver and intestine significantly reduces the adverse effects of *in vivo* reprogramming and allows stronger induction protocols^19^. To further explore the potential of this novel reprogramming strain, we used the previously generated model in which the 4F cassette was simultaneously removed from both liver and intestine using Albumin and Villin-1 Cre lines (Non Liv/Int rtTA) and changed the rtTA by the rtTA3 transactivator to enhance OSKM expression (Non Liv/Int rtTA3) (Figure 4A). Upon continuous doxycycline treatment at 1mg/ml, we observed a significant reduction in both median lifespan and body weight in Non Liv/Int rtTA3 compared to Non Liv/Int rtTA mice (Figure 4B-C). As expected, the level of *Oct4* and *Sox2* expression after 4 days of doxycycline treatment were significantly increase in Non Liv/Int rtTA3 compared to Non Liv/Int rtTA mice or 4FjF rtTA HET mice (Figure 4D and Figure S3A).

**Figure 4.**
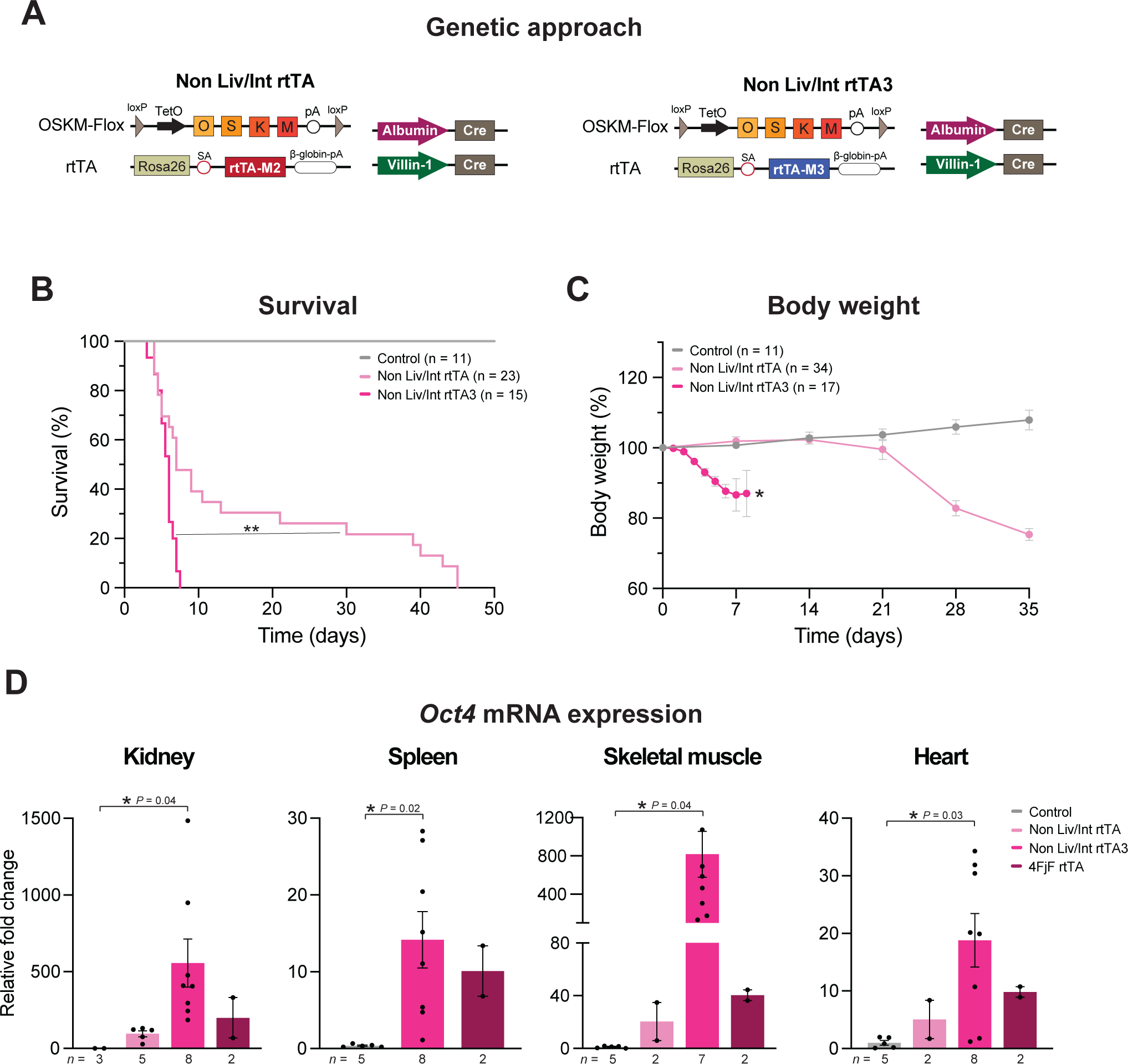
Enhanced tissue specific reprogramming avoiding the liver and the intestine. **A**, Schematic representation of the 4F Non-Liv/Int rtTA and 4F Non Liv/Int rtTA3 mouse strains carrying the polycistronic cassette for the mouse 4F between two loxP sites, OSKM-Flox, the rtTA-M2 and rtTA-M3 in *Rosa26* locus, respectively and Cre recombinase under the control of the promoter of the *Albumin* and *Villin-1* genes. **B,** Survival of control, Non Liv/Int rtTA and Non Liv/Int rtTA3 mice upon continuous administration of doxycycline. **C,** Body weight changes in control (WT), Non Liv/Int rtTA and Non Liv/Int rtTA3 mice upon continuous administration of doxycycline. **D,** *Oct4* mRNA transcript levels in multiple organs in control (WT), Non Liv/Int rtTA and Non Liv/Int rtTA3 mice after 4 days of doxycycline treatment. Data shown mean ± standard deviation. Statistical significance was assessed by one-way ANOVA followed by Tukey’s post hoc test (c,d) and log-rank (Mantel–Cox) test (b). See also Figure S3.

In general, these results demonstrate that replacing the regular rtTA transactivator with rtTA3 and avoiding the expression of OSKM in the liver and intestine increases the expression levels of the factors, however, it is still associated with adverse effects probably due to the effects of reprogramming in other tissues and organs.

### Chimeric *in vivo* reprogramming reduces adverse phenotypes and mortality

Towards the goal of achieving high and systemic induction of the reprogramming factors and reducing the adverse effects of *in vivo* reprogramming, we have generated novel reprogrammable mouse strains based on innovative genetic strategies. First, we generated a reprogrammable mouse strain in which the expression of reprogramming factors could be induced only in post-mitotic cells. Briefly, the 4FjF rtTA-M2 reprogrammable mouse (4FjF rtTA HET) was crossed with Ki67Cre^ERT2^ mice to remove the 4F cassette in actively dividing cells upon tamoxifen injection, generating 4F-Flox Ki67 mice (Figure 5A). Expression of Cre recombinase was confirmed in the intestine, but not in the liver, after a round of tamoxifen injection (Figure S4A). Based on our previous results, the number of Ki67 cells in the liver increases after reprogramming^19^, therefore, in order to prevent the adverse effect of *in vivo* reprogramming in this organ, we activated recombination after a cycle of doxycycline before tamoxifen injections (2nd) (Figure 5A). While no differences in survival were observed following the normal protocol of induction (1st), the 4F-Flox Ki67Cre mice showed a significant increase in median survival compared to whole-body mice (4FjF rtTA HET) following the second induction protocol (2nd) (Figure 5B). Consistently, the first protocol resulted in a reduction of body weight similar to that observed in 4FjF rtTA HET mice when reprogramming was induced. However, no major changes in body weight were observed in mice that followed the second induction protocol probably due to the removal of the cassette from the rapidly dividing cells in the small intestine (Figure 5C), and suggesting that at least 2x rounds of tamoxifen enhance the removal of the cassette, allowing a regular intestinal function. We then studied the *Oct4* and *Sox2* transcript levels in several tissues. Importantly, a tendency towards a reduction in the levels of OSKM was observed in the liver and small intestine of 4F-Flox Ki67 mice following the 2nd protocol compared to the whole-body 4FjF rtTA HET reprogrammable mice, while the levels of reprogrammable factors were comparable in other tissues, such as the kidney and spleen (Figure 5D and Figure S4B).

**Figure 5.**
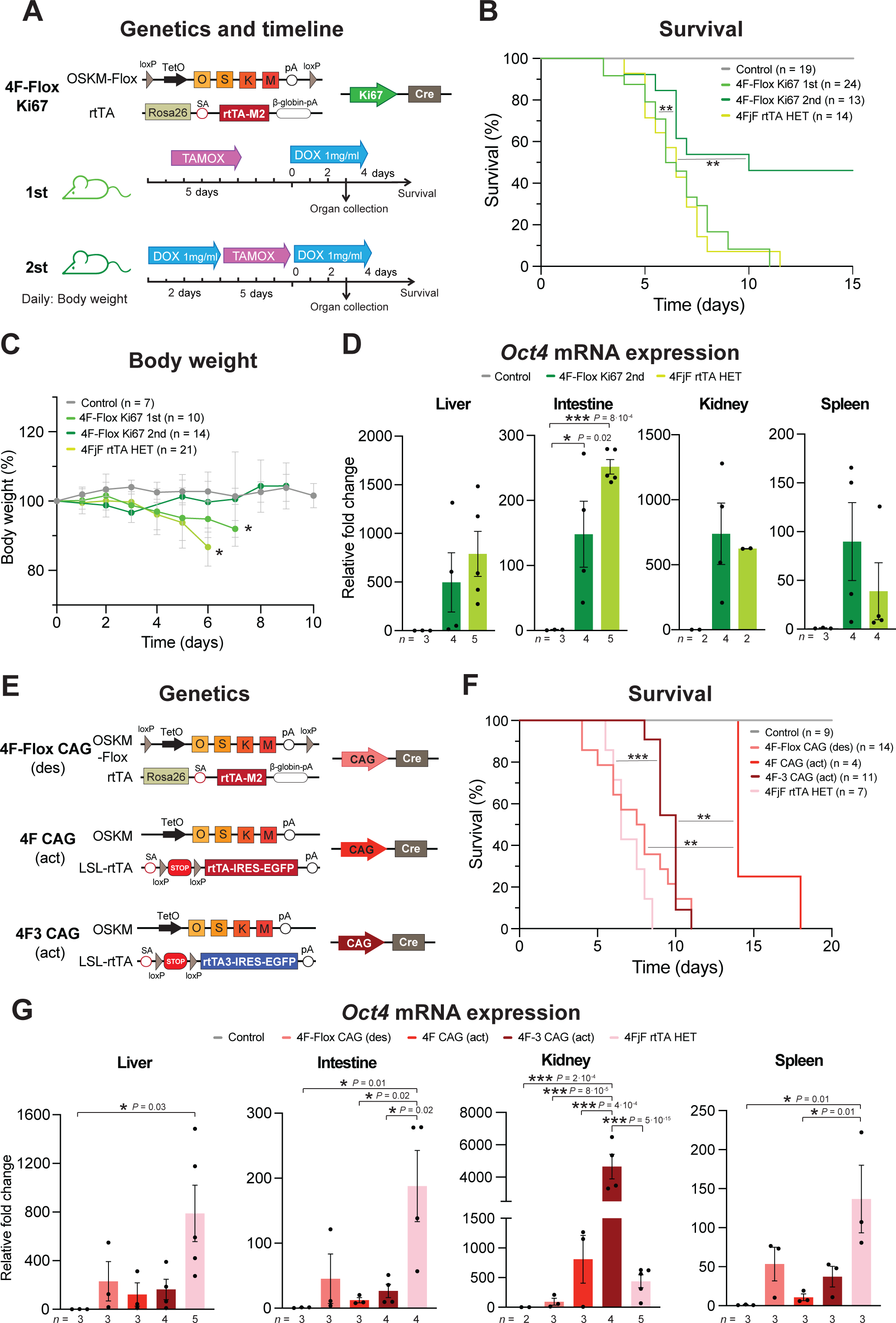
Chimeric reprogramming transgenic strains showed extended survival. **A**, Graphical representation of the 4-Flox Ki67 strain carrying the polycistronic cassette for the mouse 4F between two loxP sites, OSKM-Flox, the rtTA-M2 in *Rosa26* locus, and Cre recombinase under the controls of the promoter *Ki67.* Below are showed the two strategies followed for the inductions of reprogramming factors, including one round of continuous doxycycline after tamoxifen injections (1st) and two rounds of doxycycline (2 days and continuous) and in between the tamoxifen injections (2nd). **B,** Survival of control (WT), 4FjF and 4F-Flox Ki67mice upon continuous administration of doxycycline followed by non, or a cycle on two days of doxycycline prior to tamoxifen injections respectively. **C,** *Oct4* mRNA transcript levels in multiple organs in control (WT), 4FjF and 4F-Flox Ki67 (following the second cycle of doxycycline) mice after 4 days of doxycycline treatment. **D,** Graphical representation of the 4-Flox CAG (deactivation) strain carrying the polycistronic cassette for the mouse 4F between two loxP sites, OSKM-Flox, the rtTA-M2 in *Rosa26* locus, and Cre recombinase under the controls of the promoter *CAG.* The 4F CAG and 4F3 CAG (activation) strains carrying the polycistronic cassette for the 4F, the rtTA-M2 or rtTA-M3, respectively, in *Rosa26* locus preceded by a stop codon flanqued by two loxP sites and Cre recombinase under the controls of the promoter *CAG*. **E,** Survival of control (WT), 4F-Flox CAG, 4F CAG, 4F-3 CAG and 4FjF mice upon continuous administration of doxycycline. **F,** *Oct4* mRNA transcript levels in multiple organs in control (WT), 4F-Flox CAG, 4F CAG and 4F-3 CAG mice after 4 days of doxycycline treatment. Data shown mean ± standard deviation. Statistical significance was assessed by one-way ANOVA followed by Tukey’s post hoc test (c,f) and log-rank (Mantel–Cox) test (b,e). See also Figure S4.

Continuing with the strategy of reducing the negative effects of *in vivo* reprogramming, we generated two chimeric reprogrammable strains to activate and control the expression of reprogramming factors in a fraction of cells upon tamoxifen administration. The first strategy was based on the of crossing 4F-Flox HET reprogrammable mice with CAG-Cre^ER^ to generate 4F-Flox CAG reprogrammable mice. In the second strategy, we crossed the whole-body 4Fj LSLrtTA mice or 4Fj LSLrtTA3 with CAG-Cre^ER^ to generate 4F CAG and 4F-3 CAG reprogrammable mice, respectively. Upon tamoxifen administration, the 4F cassette (in 4F-Flox CAG mice) or the codon stop in the LSLrtTA and LSLrtTA3 transactivators (in 4F CAG and 4F-3 CAG mice) were removed, modulating the expression of reprogramming factors in CAG-positive cells (Figure 5E). Upon continuous administration of doxycycline, all the strains showed an increase in survival, with a more significant increase in median survival in the case of the activation strategy (4F CAG and 4F-3 CAG) compared to the whole-body 4FjF rtTA HET reprogrammable mice (Figure 5F). Surprisingly, survival observations where not correlated with reduction in body weight (Figure S4C). We then analyzed the *Oct4* and *Sox2* transcript levels in several tissues, observing a reduction in expression in the liver, small intestine, and spleen in all chimeric strains, compared to the 4FjF rtTA HET reprogrammable mice. On the other hand, expression in the kidney of 4F CAG and 4F-3 CAG mice was significantly increased compared to whole-body 4FjF rtTA HET mice (Figure 5G and Figure S4D).

Overall, these results demonstrate that induction of *in vivo* reprogramming in a chimeric pattern might allow to preserve organ function by reducing the number of cells expressing the factors in organs such as the liver and small intestine leading to increased survival, while simultaneously allowing a high expression of the Yamanaka factors in other tissues and organs.

## Discussion

The systematic understanding of whole-organism partial *in vivo* reprogramming in mice is grounded in the concept of inducing an epigenetic transition to a ‘youthful’ state without compromising cell identity. In recent years, many studies have shown that reprogramming can reverse age-associated features *in vivo*, therefore identifying the expression levels of factors and treatment duration as crucial components of the reprogramming process^22^. Still, reference protocols for achieving safe and potent induction of *in vivo* reprogramming have not been established. Remarkably, Abad et al. showed that a continuous induction protocol led to teratoma development and death in the 4FsB strain^16^. Subsequently, Ocampo et al. performed partial reprogramming by short-term cyclic expression of OSKM, mitigating adverse effects, improving cellular and physiological hallmarks of aging, and extending the lifespan in a mouse model of premature aging^11^. More recently, this cyclic protocol was simplified, revealing that even a single short treatment (2.5-week period) can improve long-term outlook (increased lifespan by 15%) in heterozygous progeric and non-progeric mice ^23^. Consequently, the modulation and dynamics of OSKM expression is a key variable that requires consideration in order to ensure safety and efficacy in the reprogramming process. To this extend, we decided to perform a comparative analysis of reprogramming induction strategies in multiple reprogrammable mouse strains across different tissue and organs, providing a deep resource for future *in vivo* reprogramming research.

### Importance of the genetic background

Our initial objective was to perform, for the first time, a side-by-side comparison of previously established reprogramming strains. For this reason, we decided to compare 1) the survival and changes in body weight under continuous treatment (1 mg/ml doxycycline), 2) the expression of *Oct4* and *Sox2* in different tissues with the same induction protocol (1 mg/ml doxycycline) and for the same duration (4 days), and 3) global transcriptional changes induced by *in vivo* reprogramming. As expected, decrease in survival and body weight correlated with higher expression of the reprogramming factors. Interestingly, this occurred in strains in which the transgene is inserted in the *Col1a1* locus (4Fj and 4Fk), compared to the other strains (4sA and 4sB). This suggests that the location of the transgene, as well as the order of the Yamanaka factors, can impact OSKM expression across tissues and organs and may explain differences observed in different strains^22^. In this line, teratoma formation reported by Abad et al. following long-term continuous induction in 4sB mice, which is not observed in 4Fj and 4Fk due to their early lethality, as well as the differences in gene expression between 4Fj and 4Fk strains. Regarding transcriptomic changes, we observed a correlation between the expression of OSKM and DEGs, particularly in the liver and intestine. Intriguingly, despite lower OSKM expression, we still found DEGs in tissues like the heart, brain or spleen, suggesting that the expression of OSKM in other tissues might have also cell-extrinsic effects in distant organs.

### OSKM expression can reach a threshold

Next, we decided to modulate the dynamics of OSKM expression using two genetic strategies: 1) increasing the number of copies of the cassette from heterozygosity to homozygosity, or 2) changing the type of transactivator (from rtTA to rtTA3) based on previous studies where the reprogramming efficiency *in vitro* was increased^21^. In both cases, we observed that survival and weight decreased dramatically after continuous treatment, correlating with an increase in OSKM expression. As the differences in survival and body weight are more pronounced when using the rtTA3 transactivator, this might suggest a higher expression, however, this was not the case at least in the tissues and organs that were analyzed. We hypothesized that the phenotypes associated with *in vivo* reprogramming in the rtTA3 strains results from inducing the expression in a greater number of cells or achieving the expression of OSKM in tissues and organs that are not reached in the other strains.

We hypothesize that something similar may be happening when increasing the concentration of doxycycline (1, 2, and 5 mg/ml) to induce *in vivo* reprogramming. Although we observed a reduction in survival and significant changes in body weight, the expression levels in the tissues and organs analyzed were comparable among the different concentrations. One potential explanation could be that in specific tissues where OSKM expression is already high, a threshold is reached, and additional doxycycline fails to further increase expression levels or that we achieved expression in other tissues and organs that were not analyzed.

### High expression of OSKM is still toxic in the Non-Liver/Intestine reprogrammable mice

Our previous studies have demonstrated that whole-body *in vivo* reprogramming leads to toxic effects including loss of body weight and premature death. Therefore, avoiding this toxicity could be a better strategy for improving the effects of *in vivo* reprogramming. In this line, a recent study from our laboratory^19^ demonstrated that *in vivo* reprogramming results in hepatic and intestinal dysfunction, representing a major cause of early mortality. Consequently, bypassing OSKM expression in the liver and intestine significantly mitigates these adverse effects and improves survival. Therefore, we decided to investigate the increase in expression of the reprogramming factors in these lines by substituting rtTA by rtTA3. In contrast to the observations in Parras et al., it is high likely that similar to the 4FjF rtTA3 HET strain, a high level of expression affects numerous tissues, leading to organ dysfunction and premature death (in the absence of tumors). It would be interesting to further investigate the potential causes of death in these mice to determine whether by bypassing other tissues we could achieve safer reprogramming.

### A new generation of reprogramming strains

Lastly, our results suggest that *in vivo* reprogramming leads to tissues and organ dysfunction as a primary cause of death. Therefore, we hypothesize that a strategy based on the expression of the reprogramming factors in non-dividing cells or in a certain percentage of cells within an organ might allow to achieve better reprogramming efficiencies without detrimental effects. To achieve this goal, we created two novel models of reprogrammable mice using Ki67 Cre and CAG Cre. Importantly, using this novel reprogramming strains we observed a significant increase in survival without decrease in body weight compared to whole-body reprogrammable strains. Furthermore, we have achieved less expression in organs such as liver or intestine, therefore decreasing the hepatic and small intestinal dysfunction, and increasing the expression in other organs such as kidney.

Our goal is that these novel strains represent new tools for the field to achieve stronger reprogramming protocols. Nevertheless, future studies will be necessary to determine the best induction protocols in these novel strains (such as adding more rounds of tamoxifen) to achieve safer and more efficient rejuvenation by *in vivo* reprogramming.

In summary, here we present an atlas of *in vivo* reprogramming across strains and tissues, providing a valuable tool to the field. In addition, we describe new *in vivo* reprogramming strategies and generate the next generation of reprogrammable mouse strains to continue research in this new and exciting field of research. We hope that new studies using these tools will allow a better understanding of the fundamental principles around organismal rejuvenation by *in vivo* reprogramming as well as the development of novel therapeutic approaches to improve human health at old age.

## Methods

### Animal housing

All experimental procedures were performed in accordance with Swiss legislation, after approval from the local authorities (Cantonal Veterinary Office, Canton de Vaud, Switzerland). Animals were housed in groups of five mice per cage with a 12hr light/dark cycle between 06:00 and 18:00 in a temperature-controlled environment at around 25°C and humidity between 40 and 70% (55% on average), with free access to water and food. Wild-type (WT) and transgenic mouse models used in this project were generated by breeding and maintained at the Animal Facility of Epalinges and the Animal Facility of the Department of Biomedical Science of the University of Lausanne.

### Mouse strains

All WT and transgenic mice were used on the C57BL/6J background. The whole-body reprogrammable mouse strain 4Fj rtTA-M2, carrying the OSKM polycistronic cassette inserted in the *Col1a1* locus and the rtTA-M2 trans-activator in *the Rosa 26* locus (rtTA-M2), was generated in the laboratory of Professor Rudolf Jaenisch^24^ and purchased from The Jackson Laboratory, Stock No: 011004. The reprogrammable mouse strain 4Fs-B rtTA-M2 and 4Fs-A rtTA-M2, carrying the OSKM polycistronic cassette inserted in the *Pparg* and Neto2 locus and the rtTA-M2 trans-activator in *Rosa 26* locus (rtTA-M2), were previously generated by professor Manuel Serrano ^16^ and kindly generously donated by professor Andrea Ablasser. The 4Fk rtTA-M2 carrying the OKSM polycistronic cassette inserted in the *Col1a1* locus and the rtTA-M2 trans-activator in *the Rosa 26* locus (rtTA-M2), was generated in the laboratory of Professor Konrad Hochedlinger^25^ and generously donated by him.

The 4F Non-Liv/Ins rtTA-M2 mouse strain was generated by breeding the 4F-Flox strain, carrying loxP sites flanking the 4F cassette in the *Col1a1* locus, previously generated by Professor Jaenisch ^24^ and purchased from The Jackson Laboratory, Stock No: 011001, with Albumin-Cre (Stock No 003574) and Villin-Cre ( Stock No 021504) mice to specifically remove the 4F cassette in liver and intestine.

The 4F Non-Liv/Ins rtTA-M3 mouse strain was generated by substituting the rtTA-M2 of the 4F-Flox strain for the rtTA-M3, created by the laboratory of Dr. Scott W. Lowe^26^ and purchased in The Jackson Laboratory, Stock No: 029627. The resultant offspring was crossed with Albumin-cre and Villin1-cre. The 4F-Flox Ki67 and 4F-Flox CAG strains were generated by breeding the 4F-Flox strain, with Ki67– Cre, purchased from The Jackson Laboratory, Stock No: 029803 or CAG-Cre purchased from The Jackson Laboratory, Stock No: 004682 respectively.

The 4F CAG and 4F3 CAG mouse strains were generated by substituting the rtTA-M2 and the rtTA-M3, of the 4Fj with a lox-stop-lox rtTA (LSLrtTA, purchased from The Jackson Laboratory, Stock No: 005670; and LSLrtTA3 purchased from The Jackson Laboratory, Stock No: 029633) respectively. All transgenic mice carry the mutant alleles in heterozygosity apart from 4F OSKM polycistronic cassette in the 4FjF rtTA HOM strain.

### Doxycycline administration

*In vivo* expression of OSKM in all reprogrammable mouse strains was induced by continuous or 3-4 days administration of doxycycline (Sigma, D9891) in drinking water (1 mg/ml, 2mg/ml, or 5mg/ml) in 2-3-month-old mice.

### Tamoxifen injections

To activate Cre-mediated recombination, intraperitoneal tamoxifen injections were administrated for five consecutive days (20mg/ml dissolved in 10% ethanol – 90% sunflower oil) at 67mg/Kg/day.

### Mouse monitoring and euthanasia

All mice were monitored at least three times per week. Upon induction of *in vivo* reprogramming, mice were monitored daily to evaluate their activity, posture, alertness, body weight, presence of tumors or wound, and surface temperature. Mice were euthanized according to the criteria established in the scoresheet. We defined lack of movement and alertness, presence of visible tumors larger than 1cm^3^ or opened wounds, body weight loss of over 30% and surface temperature lower of 34°C as imminent death points. For survival, body weight experiments as well as tissue and organ collection, mice of both genders were randomly assigned to control and experimental groups. Animals were sacrificed by CO_2_ inhalation (6 min, flow rate 20% volume/min). Subsequently, animals were perfused the mice with saline. Finally, multiple organs and tissues were collected in liquid nitrogen and used for DNA or RNA extraction.

### RNA extraction

Total RNA was extracted from mouse tissues and organs using TRIzol (Invitrogen, 15596018). Briefly, 500 µl of TRIzol was added to 20-30 µg of frozen tissue into a tube (Fisherbrand 2 ml 1.4 Ceramic, Cat 15555799) and homogenized at 7000 g for 1 min using a MagNA Lyser (Roche diagnostic) at 4°C. Subsequently, 200 µl of chloroform was added to the samples and samples were vortexed for 10 sec and placed on ice for 15 min. Next, samples were centrifuged for 15 min at 12000 rpm at 4°C and supernatants were transferred into a 1.5 ml vial with 200 µl of 100% ethanol. Finally, RNA extraction was performed using the Monarch total RNA Miniprep Kit (NEB, T2010S) following the manufacture recommendations and RNA samples were stored at −80°C until use.

### cDNA synthesis

Total RNA concentration was determined using the Qubit RNA BR Assay Kit (Q10211, Thermofisher), following the manufacture instructions and a Qubit Flex Fluorometer (Thermofisher). Prior to cDNA synthesis, 2 µL of DNAse (1:3 in DNase buffer) (Biorad, 10042051) was added to 700 ng of RNA sample, and then incubated for 5 min at room temperature (RT) followed by an incubation for 5 min at 75°C to inactivate the enzyme. For cDNA synthesis, 4 µL of iScript™ gDNA Clear cDNA Synthesis (Biorad, 1725035BUN) was added to each sample, and then placed in a thermocycler (Biorad, 1861086) following the following protocol: 5 min at 25°C for priming, 20 min at 46°C for the reverse transcription, and 1 min at 95°C for enzyme inactivation. Finally, cDNA was diluted using autoclaved water at a ratio of 1:5 and stored at −20°C until use.

### RNA-seq alignment and quantification

Data was processed using nf-core/rnaseq v3.14.0 (doi: https://doi.org/10.5281/zenodo.1400710) of the nf-core collection of workflows^27^, using reproducible software environments from the Bioconda^28^ and Biocontainers^29^ projects. The pipeline was executed with Nextflow v23.10.0^30^ with the following command: RNA-seq reads were aligned to the *Mus musculus* reference genome GRCm39 (GCA_000001635.9), and gene annotation was obtained from the Ensembl release 110.

### RNA-seq analysis

Transcript read counts were imported into R and converted to gene counts using the Bioconductor package ‘tximport’^31^. Normalization was conducted using the Bioconductor package ‘DESeq2’ [Love MI], and dimensionality reduction was performed using the reads counts after variance stabilizing transformation (VST). Differential expression analysis was carried out using the ‘DESeq2’ package with the parameter modelMatrixType set to “standard” and default settings for other parameters. Genes were considered differentially expressed between conditions if they exhibited an adjusted p-value below 0.05 and an absolute log2 fold change exceeding 2. Gene ontology analysis was conducted using the ‘compareCluster’ function from the Bioconductor package ‘clusterProfiler’ [Wu T]. The analysis utilized the org.Mm.eg.db database^32^, focusing solely on Biological Process (BP) ontology. Benjamini-Hochberg adjustment was applied for p-values, with significance thresholds set to 0.05 for both p-values and q-values.

### Semiquantitative RT-PCR

The following specific primers were used to detect the expression of the Cre recombinase in the cDNA of mice samples, Cre forward: 5’-GAACGAAAACGCTGGTTAGC-3’, and Cre: reverse 5’-CCCGGCAAAACAGGTAGTTA-3’ at a final concentration of 0.4 µM. DNA was amplified using DreamTaq Green PCR Master Mix 2X (Thermofisher, K1081) following the amplification protocol: 3 min at 95°C + 33 cycles (30 s at 95°C + 30 s at 60 °C + 1 min at 72°C) + 5 min at 72°C. PCR product were loaded and run in an agarose (1.6%) gel containing ethidium bromide (Carlroth, 2218.1). Images were scanned with a gel imaging system (Genetic, FastGene FAS-DIGI PRO, GP-07LED).

### qRT-PCR

qRT-PCR was performed using SsoAdvanced SYBR Green Supermix (Bio-Rad, 1725274) in a PCR plate 384-well (Thermofisher, AB1384) and using a Quantstudio 12K Flex Real-time PCR System instrument (Thermofisher). Forward and reverse primers were used at a ratio 1:1 and final concentration of 5 µM with 1ul of cDNA. *Oct4* and *Sox2* mRNA levels were determined using the following primers: *Oct4* forward: 5’-GGCTTCAGACTTCGCCTTCT-3’ *Oct4* reverse: 5’-TGGAAGCTTAGCCAGGTTCG-3’, *Sox2* forward: 5’-TTTGTCCGAGACCGAGAAGC-3’, *Sox2* reverse: 5’-CTCCGGGAAGCGTGTACTTA-3’. mRNA levels were normalized using the house keeping gene *Gapdh (*forward: 5’-GGCAAATTCAACGGCACAGT-3’, reverse: 5’-GTCTCGCTCCTGGAAGATGG-3’).

### Data analysis

Statistical analysis was performed using GraphPad Prism 9.4.1 (GraphPad Software). The normality of the data was studied by Shapiro-Wilk test and homogeneity of variance by Levene test. For comparison of two independent groups, two-tail unpaired t-Student’s test (data with normal distribution), Mann-Whitney-Wilcoxon or Kolmogorov-Smirnov tests (with non-normal distribution) was executed. For multiple comparisons, data with a normal distribution were analyzed by one way-ANOVA test followed by a Tukey’s (equal variances) or a Games-Howell’s (not assumed equal variances) post-hoc tests. Statistical significance of non-parametric data for multiple comparisons was determined by Kruskal-Wallis one-way ANOVA test.

## Author contributions

S.P. and A.V.-A. were involved in the design of the study, performing the experiments, data collection and statistical analysis. J.A.d-S and F.v-M were involved in RNAseq analysis. A.P. helped generating mice strains and samples. G.D., C.M., M.C.M., and C.Y.M. contributed to RNA and protein extraction, qRT–PCR and western blot analysis. C.V.B. was involved in genotyping and sample collection. A.O. directed and supervised the study and designed the experiments. S.P., A.V.-A. and A.O wrote the manuscript with input from all authors.

## Competing interests

A.O. is co-founder and shareholder of EPITERNA SA (non-financial interests) and Longevity Consultancy Group (non-financial interests). The rest of the authors declare no competing interests.

## Acknowledgements

The authors thank all members of the Ocampo laboratory for helpful discussions and continuous support. We thank the teams of mouse facilities at the University of Lausanne including Francis Derouet (head of the animal facility at Epalinges), I. Grandjean (head of the animal facility of Agora) and L. Lecomte (head of the animal facility of the Department of Biomedical Sciences). We thank K. Hochedlinger for the kind donation of the 4Fk mice. We thank M. Serrano and A. Ablasser for the kind donation of the 4Fs-A rtTA and 4Fs-B rtTA mice. This work was supported by the Milky Way Research Foundation (MWRF), the Eccellenza grants from the Swiss National Science Foundation (SNSF), the University of Lausanne, and the Canton Vaud. G.D.-M. was supported by the EMBO postdoctoral fellowship (EMBO ALTF 444-2021 to G.D.-M.).

## Funding

This work was supported by the Milky Way Research Foundation (MWRF), the Eccellenza grants from the Swiss National Science Foundation (SNSF), the University of Lausanne, and the Canton Vaud.

---

**Figure S1.**
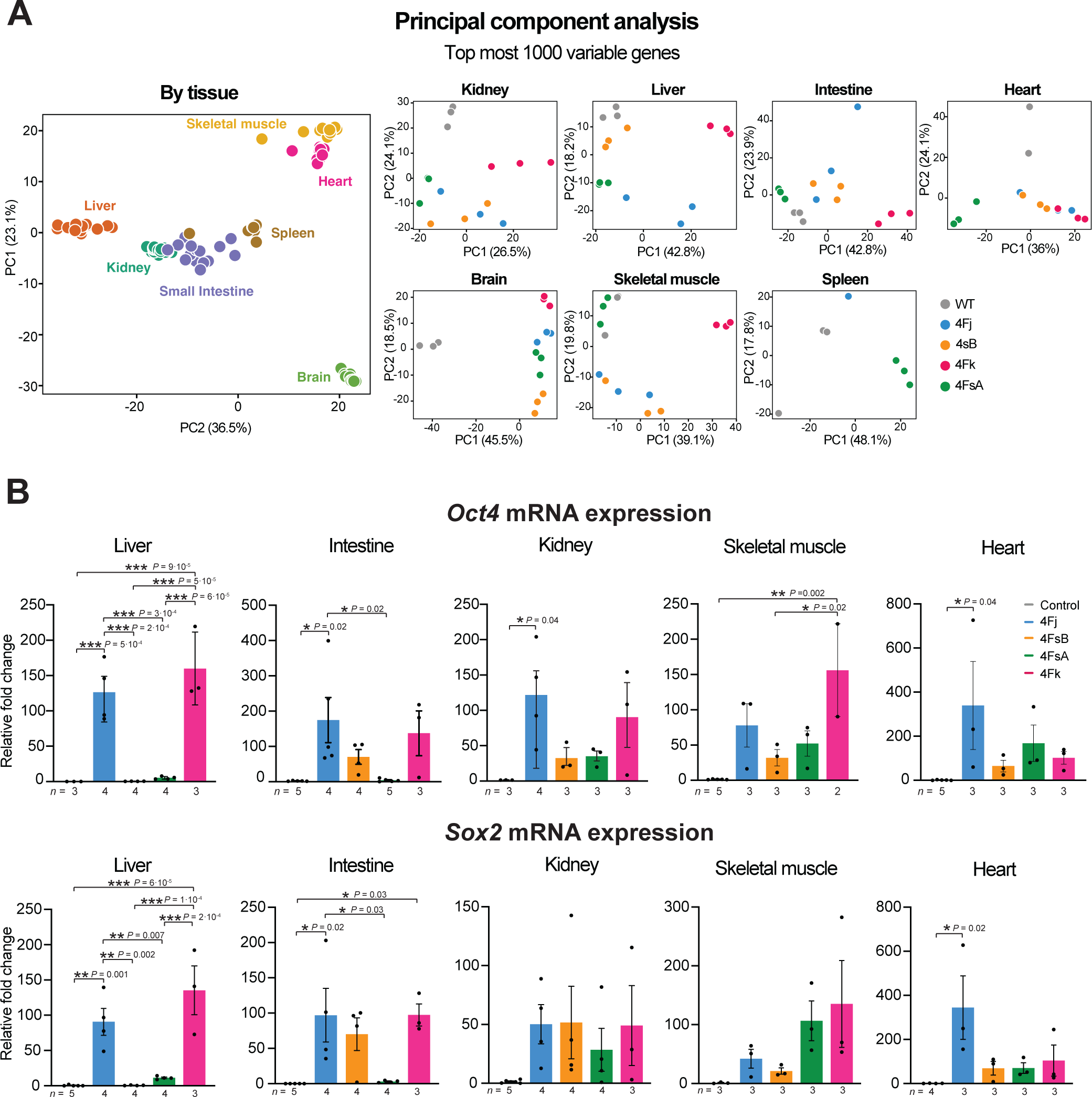
*In vivo* reprogramming in different whole body reprogrammable mouse strains. **A**, Principal component analysis (PCA) of RNA-seq data by tissue (left) and by strain (right). **B,** *Oct4 and Sox2* mRNA transcript levels in multiple organs in control (WT), 4Fj, 4FsA, 4FsB and 4Fk mice after 4 days of doxycycline treatment. Data shown mean ± standard deviation. Statistical significance was assessed by one-way ANOVA followed by Tukey’s post hoc test (b).

**Figure S2.**
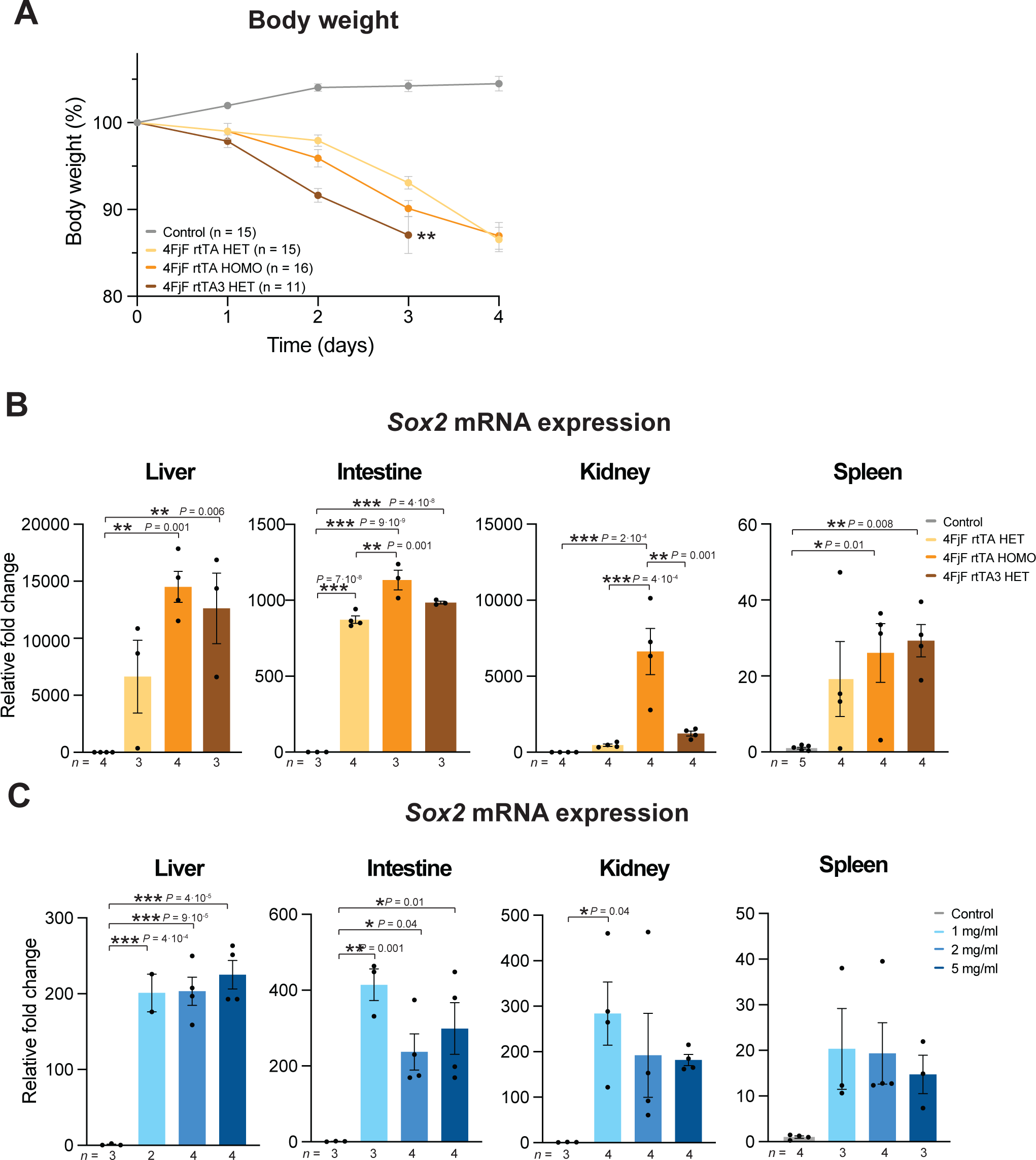
Strategies to improve *in vivo* reprogramming in whole-body reprogrammable strains. **A**, Body weight changes in control (WT), 4FjF rtTA HET, 4FjF rtTA HOMO and 4FjF rtTA3 HET mice upon continuous administration of doxycycline **B,** *Sox2* mRNA transcript levels in multiple organs in control (WT), 4FjF rtTA HET, 4FjF rtTA HOMO and 4FjF rtTA3 HET after 4 days of doxycycline treatment. **C,** *Sox2* mRNA transcript levels in multiple organs in control (WT) and 4FjF rtTA HET after 4 days of doxycycline treatment at 1mg/ml 2 mg/ml and 5mg/ml. Data shown mean ± standard deviation. Statistical significance was assessed by one-way ANOVA followed by Tukey’s post hoc test.

**Figure S3.**
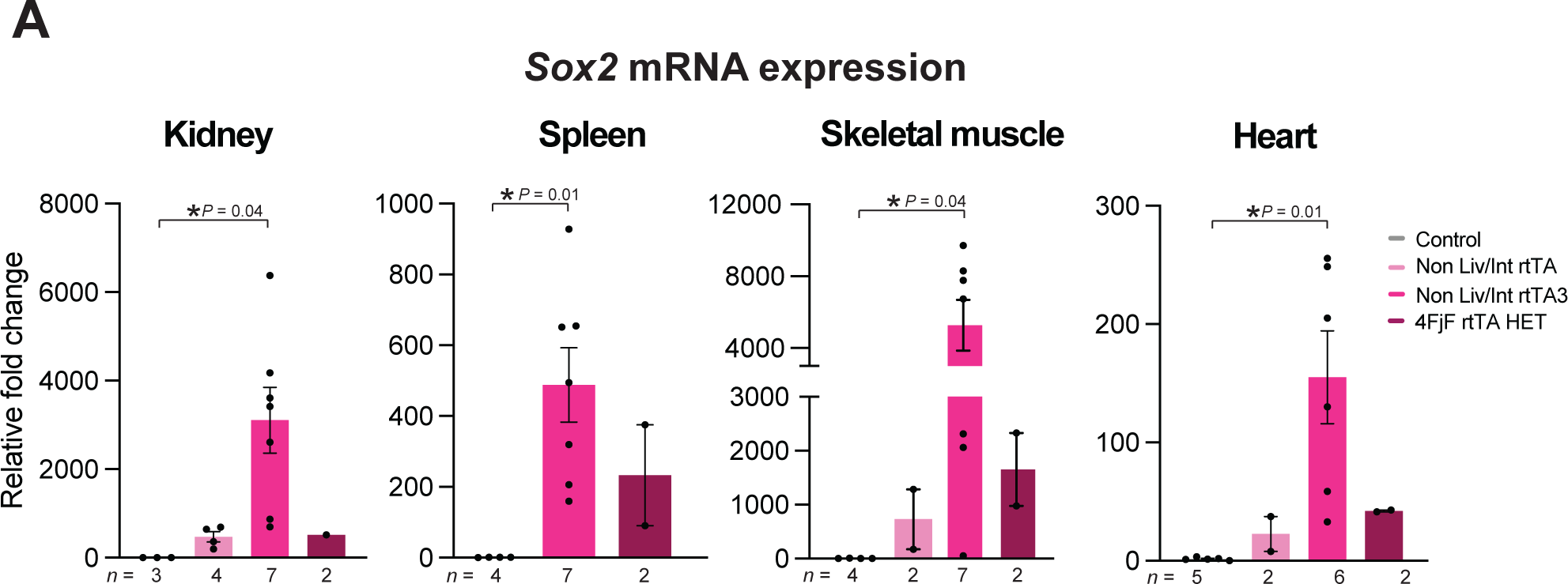
Comparative *in vivo* reprogramming in Non Liv/Int strains. **A**, *Sox2* mRNA transcript levels in multiple organs in control, Non Liv/Int rtTA and Non Liv/Int rtTA3 mice after 4 days of doxycycline treatment. Data shown mean ± standard deviation. Statistical significance was assessed by one-way ANOVA followed by Tukey’s post hoc test.

**Figure S4.**
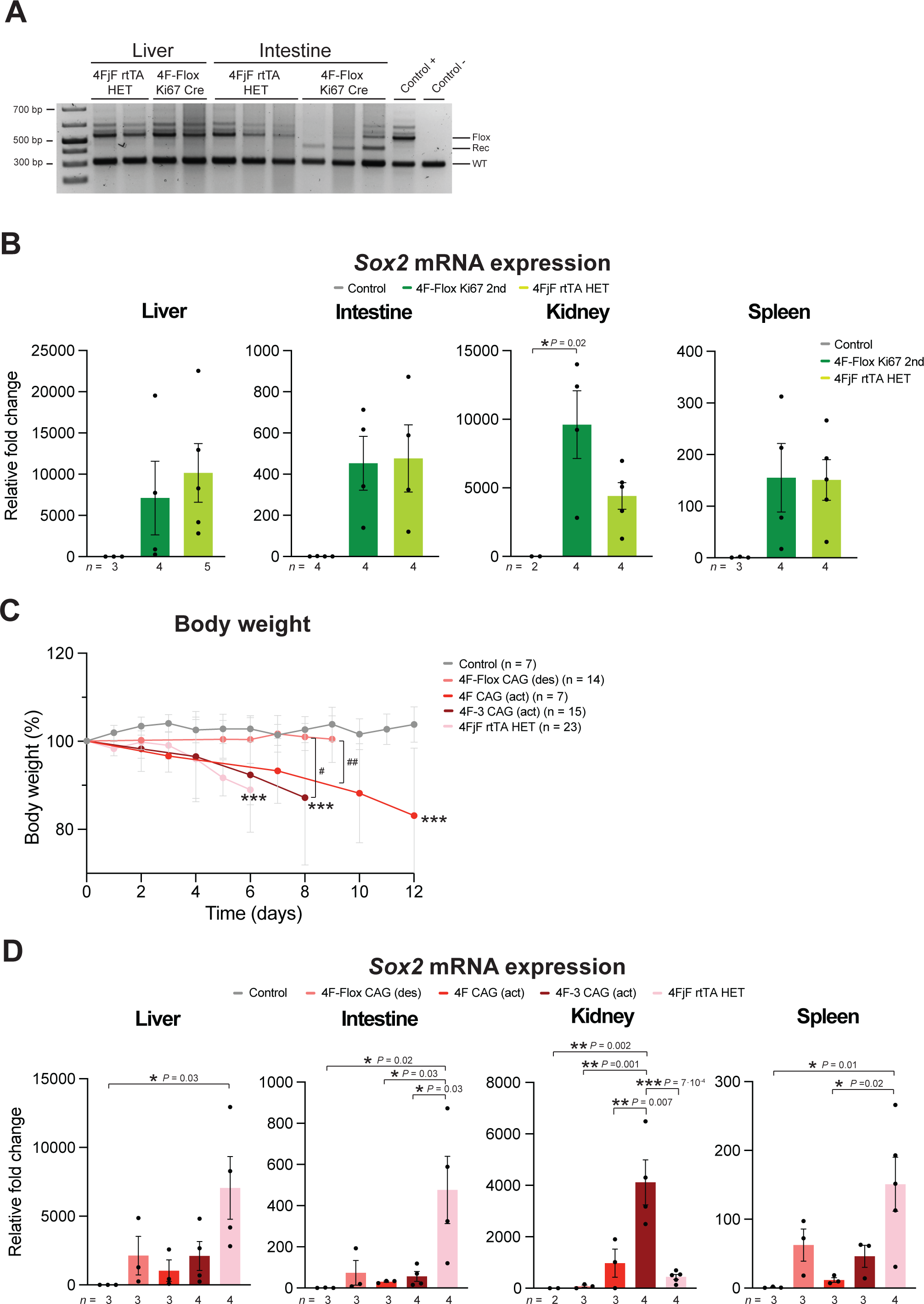
Generation of *in vivo* reprogramming chimeric strains. **A**, Expression of Cre recombinase in the liver and the small intestine of 4FjF rtTA and 4F-Flox Ki67 mice after tamoxifen injections. **B,** Body weight changes in control (WT), 4FjF and 4F-Flox Ki67 mice upon continuous administration of doxycycline followed by non, or a cycle on two days of doxycycline prior to tamoxifen injections respectively. **C,** *Sox2* mRNA transcript levels in multiple organs in control (WT), 4FjF and 4F-Flox Ki67 (following the second cycle of doxycycline) mice after 4 days of doxycycline treatment. **D,** Body weight changes of control (WT), 4F-Flox CAG, 4F CAG, 4F-3 CAG and 4FjF mice upon continuous administration of doxycycline. **e,** *Sox2* mRNA transcript levels in multiple organs in control (WT), 4F-Flox CAG, 4F CAG and 4F-3 CAG mice after 4 days of doxycycline treatment. Data shown mean ± standard deviation. Statistical significance was assessed by one-way ANOVA followed by Tukey’s post hoc test (b-e).

